# High-accuracy alignment ensembles enable unbiased assessments of sequence homology and phylogeny

**DOI:** 10.1101/2021.06.20.449169

**Authors:** Robert C. Edgar

**Affiliations:** Unaffiliated

## Abstract

Multiple sequence alignments (MSAs) are widely used to infer evolutionary relationships, enabling inferences of structure, function, and phylogeny. Standard practice is to construct one MSA by some preferred method and use it in further analysis; however, undetected MSA bias can be problematic. I describe Muscle5, a novel algorithm which constructs an ensemble of high-accuracy MSAs with diverse biases by perturbing a hidden Markov model and permuting its guide tree. Confidence in an inference is assessed as the fraction of the ensemble which supports it. Applied to phylogenetic tree estimation, I show that ensembles can confidently resolve topologies with low bootstrap according to standard methods, and conversely that some topologies with high bootstraps are incorrect. Applied to the phylogeny of RNA viruses, ensemble analysis shows that recently adopted taxonomic phyla are probably polyphyletic. Ensemble analysis can improve confidence assessment in any inference from an MSA.

## Introduction

### Background

Multiple sequence alignment (MSA) algorithms are ubiquitous in molecular biology, with popular software such as `Clustal-Omega` [1], `MAFFT` [2] and `MUSCLE` [3] receiving hundreds of citations per year. Despite decades of research into automated alignment, current algorithms predict > 30% columns incorrectly on structure-based benchmarks [4, 5]. Most alignment algorithms are based on highly simplified models of evolution parameterised by substitution scores and gap penalties. Default values for model parameters are somewhat arbitrary as they are trained on data of varying relevance to a particular set of sequences in practice. Systematic changes, and hence the opportunity for systematic errors, may be induced by changing parameters. For example, reducing gap penalties tends to increase the number of gaps. Most algorithms, including Clustal-Omega, MAFFT and MUSCLE, use progressive alignment according to a guide tree [6] which may cause bias towards the this tree e.g. in an estimated phylogeny [7]. However, standard practice is to construct a single MSA using some preferred method and proceed on the assumption that bias (henceforth understood to include alignment errors of any kind which may affect downstream inference) can be neglected.

### Muscle5

The `Muscle5` algorithm constructs a collection (*H-ensemble*) of high-accuracy MSAs (*replicates*) such that no particular MSA from the collection (or by any other method) is preferred *a priori* (see Table 1 for summary of terminology). An MSA is built following the strategy pioneered by Probcons [8]: posterior probabilities for aligning all letter pairs are computed using a hidden Markov model (HMM), a consistency transformation [9] is applied, and the final MSA is constructed by maximum expected accuracy pair-wise alignments [10] by progressive alignment according to a guide tree. HMM parameters and its guide tree are fixed in one replicate and varied between replicates to maximise differences in bias between replicates without degrading accuracy. If each replicate has different bias, then averaging results over replicates can correct for bias, and comparing results from different replicates can assess whether bias is important in a particular downstream analysis. Variations are introduced by multiplying HMM probabilities by a random number in the range −0.25 … + 0.25, the largest range found to maintain accuracy on structural benchmarks. Varying HMM probabilities is equivalent to varying gap penalties and substitution scores. The guide tree, an important potential source of bias, is also varied by permuting the joining order of large subgroups close to the root. A divide-and-conquer strategy enables scaling to tens of thousands of sequences (details and survey of related prior work in Methods and Supplementary Material).

**Table 1:**
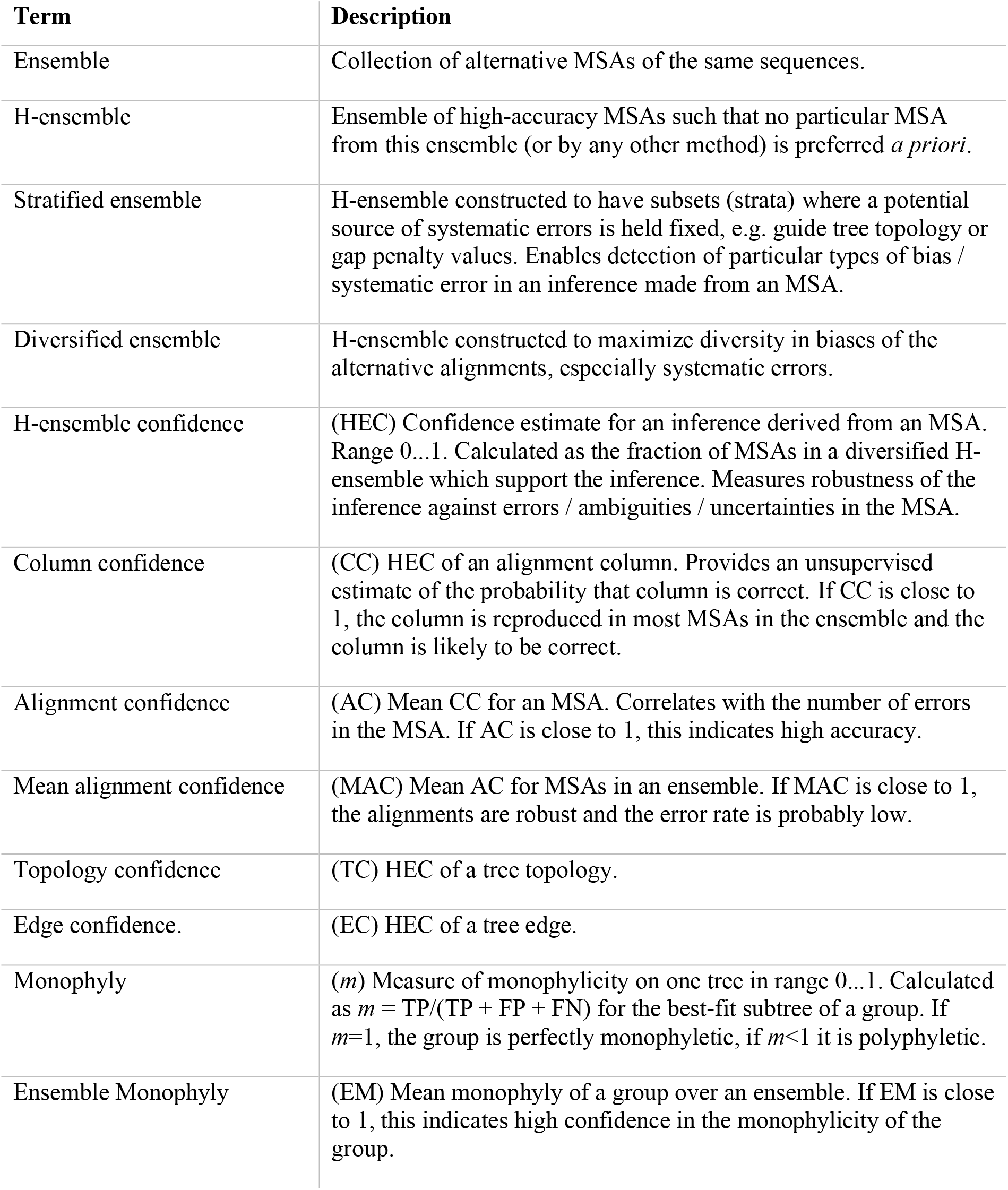
Summary of technical terms.

### Ensemble confidence measures

The *H-ensemble confidence* (HEC) of an inference from an MSA is the fraction of replicates which supports it. For example, if all replicates support the same inference then HEC=1. HEC is calculated using a *diversified* H-ensemble which is designed to generate the greatest possible variety in the alignments, especially in systematic errors, so that averaging over the ensemble mitigates MSA bias. This approach can be applied to the alignments themselves: the Column Confidence (CC) is the HEC of an alignment column, i.e. the fraction of replicates where the column is reproduced. Unlike typical conservation-based metrics, a column with many gaps or with biochemically dissimilar amino acids will be assigned high CC if it is consistently reproduced. Alignment Confidence (AC) is the mean column confidence of a replicate, and MAC is the mean AC over the ensemble. If MAC=1, all replicates are identical and the alignment is robust; if MAC is smaller then the alignment is more sensitive to small parameter adjustments. Differences between alignments necessarily reflect errors, and lower MAC values therefore necessarily indicate higher error rates in a typical MSA from the ensemble. The robustness of a phylogenetic tree against MSA bias can be assessed by comparing replicate trees, i.e. trees estimated from different alignment replicates. Edge Confidence (EC) is the HEC of a tree edge, i.e. the fraction of replicate trees where the edge is reproduced. Topology Confidence (TC) is the fraction of replicates supporting the branching order of designated subgroups such as taxonomic clades, and Ensemble Monophyly (EM) is the mean monophylicity for a subgroup. These phylogenetic tests are independent of bootstrapping as there is no re-sampling of columns.

### MSA bias detection

Guide tree bias can be assessed by comparing results on replicates where the guide tree is held fixed. If inferences differ and correlate with the choice of guide tree, then guide tree bias is present by definition. In general, a *stratified ensemble* has subsets (*strata*) where some parameters are held fixed while others are varied. Comparing results from different strata enables detection of particular types of bias in any inference from an MSA.

## Results

### Alignment accuracy

As shown in Fig. 1, the accuracy of `Muscle5` on structure-based benchmarks is higher than the state of the art represented by `Clustal-Omega` and `MAFFT`. There is negligible difference in average accuracy between parameter variants, showing that all replicates are equally plausible *a priori.* On a benchmark with 10,000 protein sequences per set, `Muscle5` aligns 59% of columns correctly, which is a 13% improvement over `Clustal-Omega` (52% columns correct) and 26% over `MAFFT` (47% correct) (Supplementary Material). Fig. 1 also shows that CC and AC provide predictive unsupervised estimates of accuracy (i.e., the estimates are independent of a trusted reference alignment). This is illustrated in Fig. 2 which shows replicate alignments of four proteins, focusing on a region with well-conserved sequence and secondary structure and a more variable surface loop where neither sequence nor structure aligns well. This transition in secondary structure is reflected in the column confidence values, where the first 15 columns have *CC* = 1.0 and later values drop to CC ≈ 0.5 in the loop, thereby identifying a segment where the alignment is error-prone.

**Figure 1:**
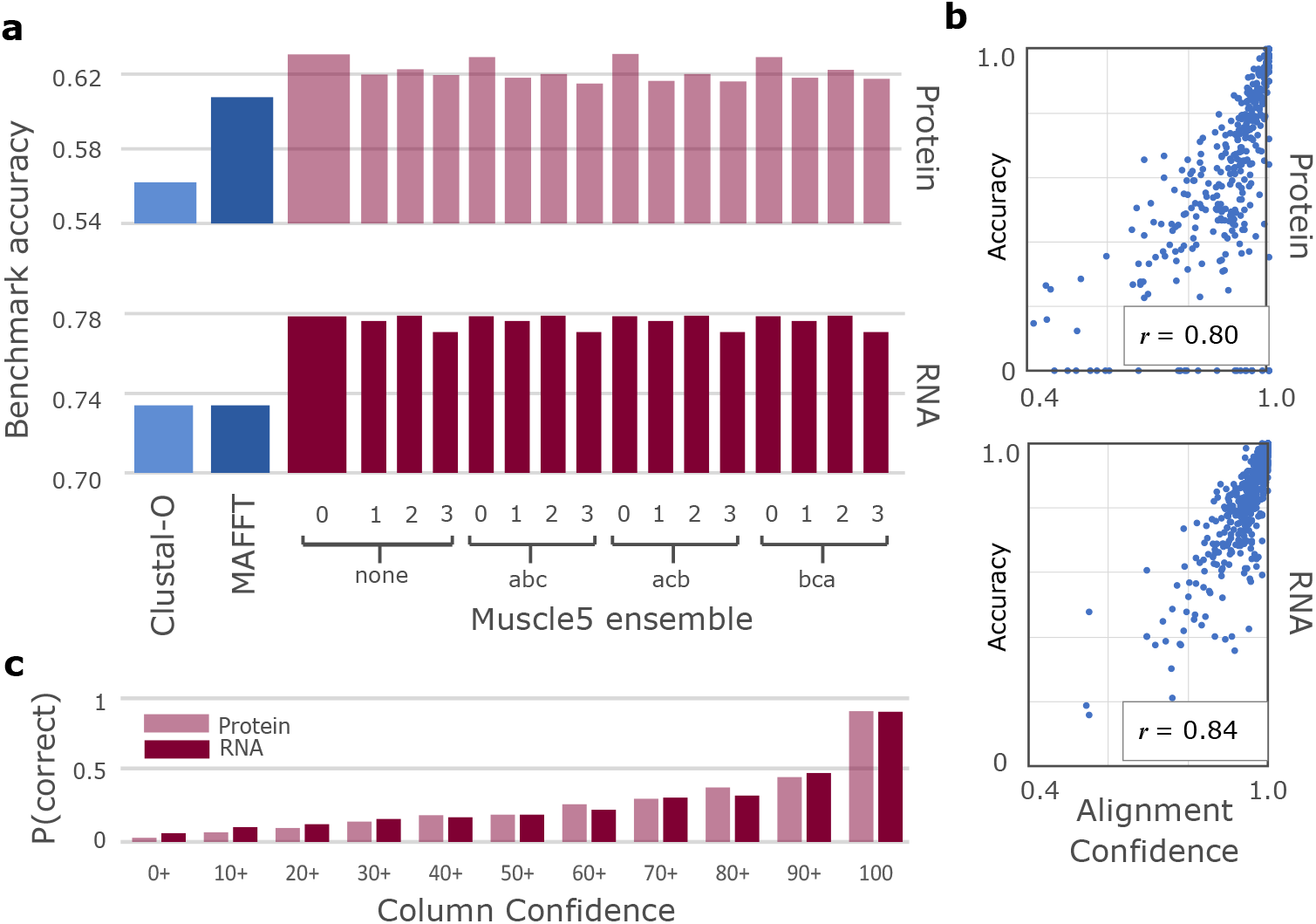
Accuracy of Muscle5 on structure-based benchmarks. **a** Average accuracy of `Muscle5` ensemble replicates compared to `Clustal-Omega` and `MAFFT` on benchmarks of protein [4] (top) and RNA [5] (bottom) alignments; the default variant *none.0* is the wider bar. **b** Correlation between AC and accuracy (fraction correct columns) for protein (top) and RNA (bottom); Pearson’s *r* = 0.80, 0.84 respectively. **c** Probability that a column is correct after binning into CC percentage intervals: 0+ is 0% ≤ *CC* < 10%, 10+ is 10% ≤ *CC* < 20% etc.; the last bar is *CC* = 100%. Thus CC is predictive of correctness and AC is predictive of accuracy.

**Figure 2:**
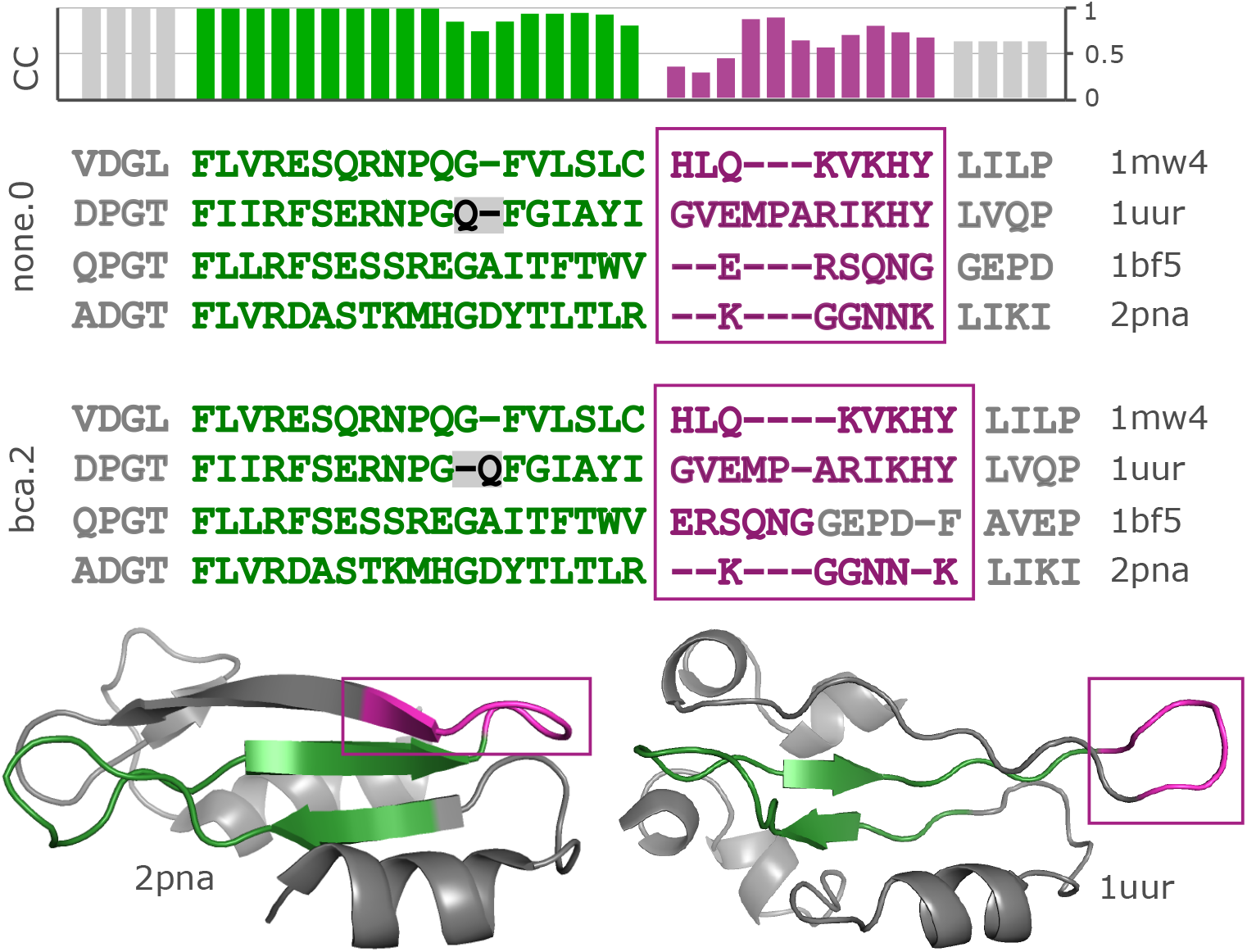
Replicate alignments of BBS11008. Two replicate alignments of a segment in `Balibase` set `BBS11008` are shown together with ribbon diagrams of two of its four structures (`2pna` and `1uur`). This region comprises a well-conserved anti-parallel beta sheet (green) which transitions into a variable exposed loop (magenta, outlined by rectangles). Sequence homology in the beta sheet is unambiguous except for one gap (grey background), while both sequence and structure similarity are unclear in the loop, which is reflected by lower CC values (top histogram; CC was calculated from a diversified ensemble of 100 replicates).

### RNA virus phylogeny

I investigated whether reported phylogenies of RNA viruses from the recent literature are reproducible and supportable, focusing on the topology of the four *Coronaviridae* genera and five *Riboviria* phyla inferred from alignments of the RNA-dependent RNA polymerase (RdRp) gene, which is widely used for phylogenetic and taxonomic analysis of viruses [11]. Coronavirus genera have well-conserved RdRp alignments with amino acid (a.a.) identities ~60%. *Riboviria* overall are highly diverged with RdRp identities often as low as 5% to 10%, representing a very challenging case. I created diversified ensembles using `Muscle5`, finding MAC was 0.91 for genus, indicating generally high confidence with some variability in the alignments, but only 0.18 for phylum, indicating a high error rate. Trees were estimated by six different methods: RAxML [12], PhyML [13], IQ-Tree [14], FastTree [15], and neighbour-joining (NJ) and minimum-evolution (ME) using MEGA [16]. Trees were rooted using outgroups *Torovirus* for genus and reverse transcriptases for phylum. As shown in Fig. 3, four of the tree methods report a strong consensus (((A,B),G),D) for the genus topology (A = *Alphacoronavirus,* B = *Betacoronavirus,* G = *Gammacoronavirus* and D = *Deltacoronavirus*), while the faster but more more approximate methods MEGA-NJ and FastTree reported more variants. The combined ensemble topology confidence of the consensus is 98.4% (82.5%) including (excluding) the latter two methods. A conventional analysis using the default `Muscle5` MSA gave low bootstrap confidence to most edges (Fig. 4). Here, ensemble analysis confidently resolves topology while a single MSA with bootstrapping does not. For phylum, there is no consensus; all tree methods report varying topologies across the ensemble (Fig. 5). Fig. 6 shows trees and bootstraps obtained on MSAs with two different guide trees and other parameters fixed. All six tree methods agree with each other on the topology according to one of these MSAs, but the topologies conflict and therefore one or both must be wrong. Further, the bootstraps of most methods are high, with values ≥ 84 for all edges from all maximum-likelihood methods except for FastTree on edge *k.* Thus, for phylum, high bootstraps for at least one incorrect topology are necessarily induced by MSA bias, and ensemble analysis shows that the topology cannot be reliably resolved. Coronavirus trees reported in recent literature are shown in Fig. 7, which are seen to have conflicting genus topologies with high bootstraps, suggesting that systematic alignment errors induced overconfidence.

**Figure 3:**
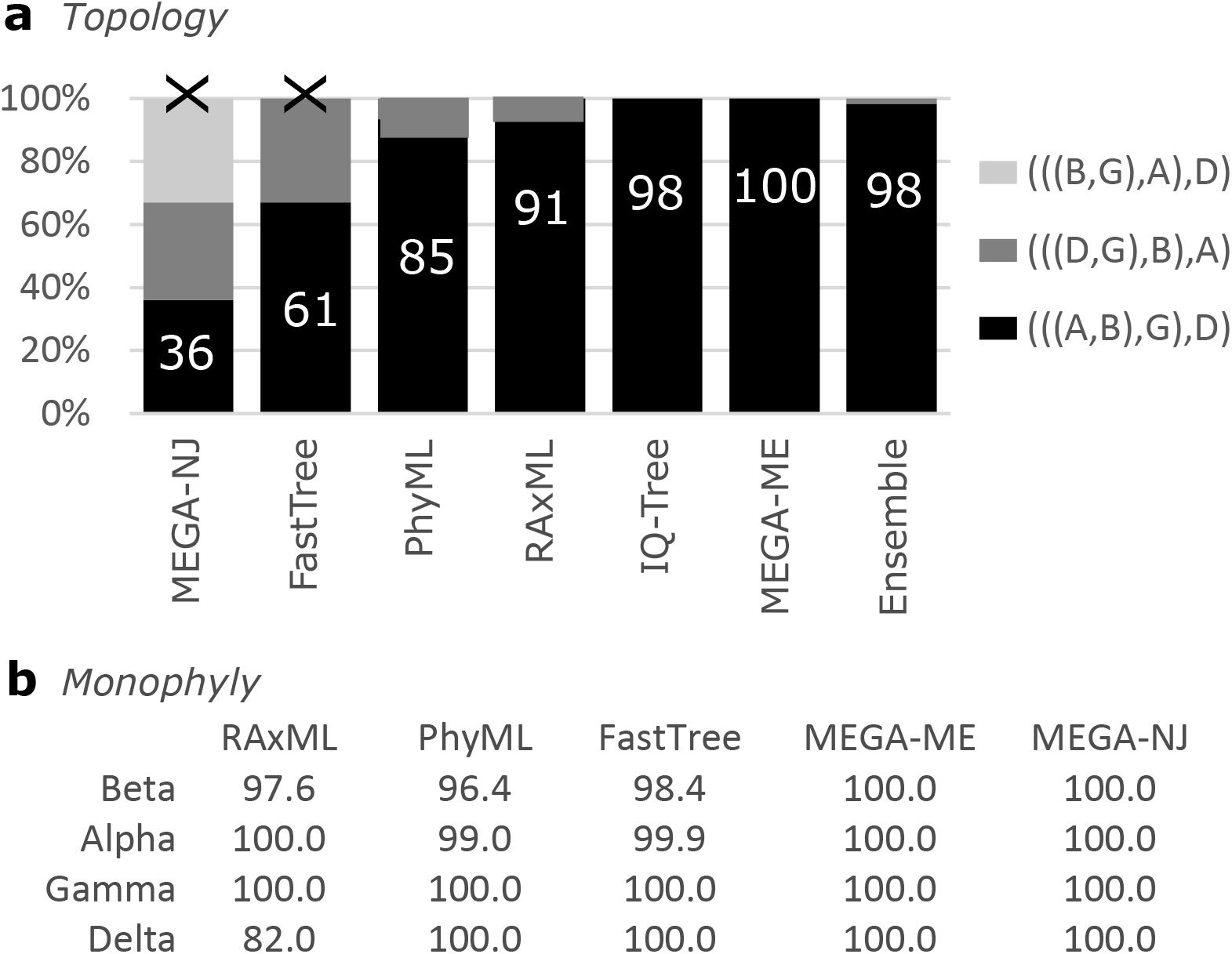
Ensemble confidence of coronavirus genus topologies and monophyly. **a** Relative frequencies of tree topologies for coronavirus genera from a diversified ensemble using six different tree estimation methods.The rightmost bar shows the combined ensemble average with MEGA-NJ and FastTree excluded. A = *Alphacoronavirus,* B = *Betacoronavirus,* G = *Gammacoronavirus* and D = *Deltacoronavirus* **b** Ensemble Monophyly (EM) of coronavirus genera. All six tree estimation methods give confidence > 96% to monophyly of all genera except for *EM* = 82.0% for Deltacoronavirus by RaxML.

**Figure 4:**
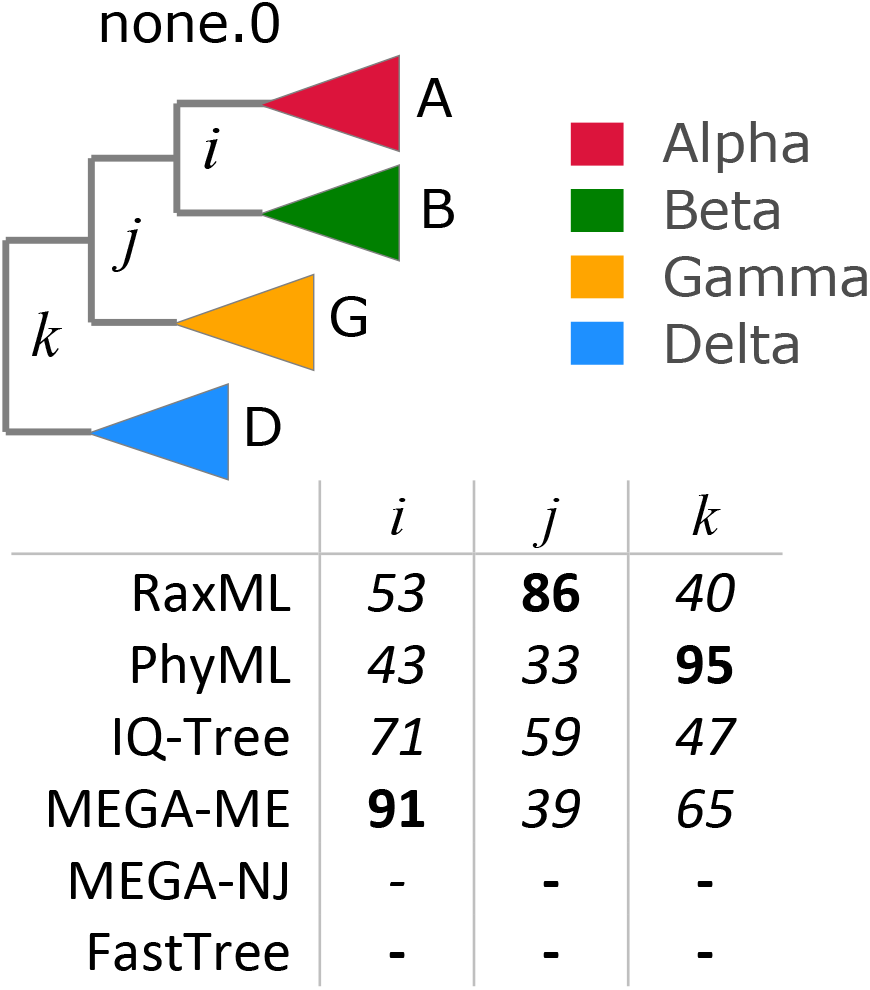
Bootstraps for coronavirus consensus genus topology. The coronavirus genus topology is `(((A,B),G),D)` with high ensemble confidence (see Fig. 3). Using the default `Muscle5` MSA (none.0) this topology was reported by four of the six tree estimation methods with bootstraps as shown in the figure, where bootstrap values are mostly low.

**Figure 5:**
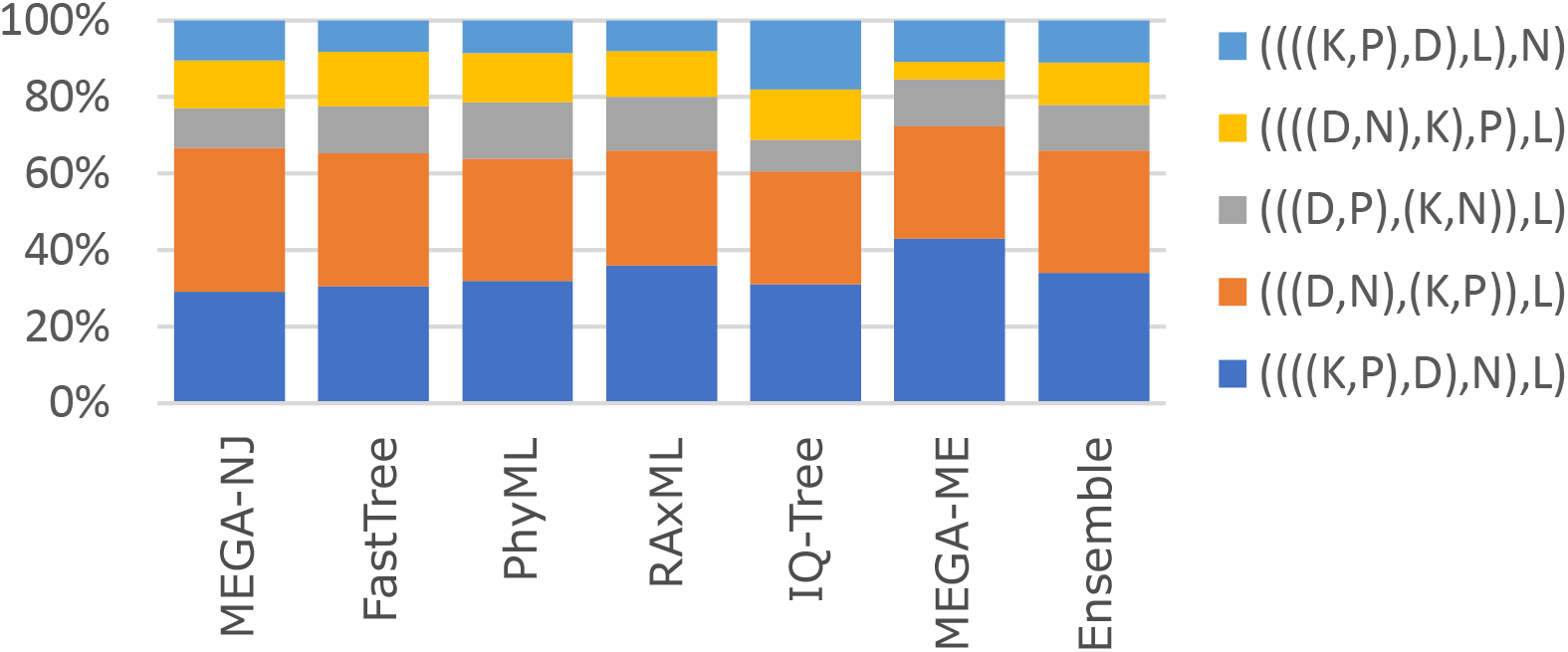
Ensemble frequencies of phylum topologies. Relative frequencies of tree topologies of Ribovirus phyla from a diversified ensemble using six different tree estimation methods. The rightmost bar shows the combined ensemble average. D = *Duplornaviricota*, K = *Kitrinoviricota,* L = *Lenarviricota,* N = *Negarnaviricota,* P = *Pisuviricota.*

**Figure 6:**
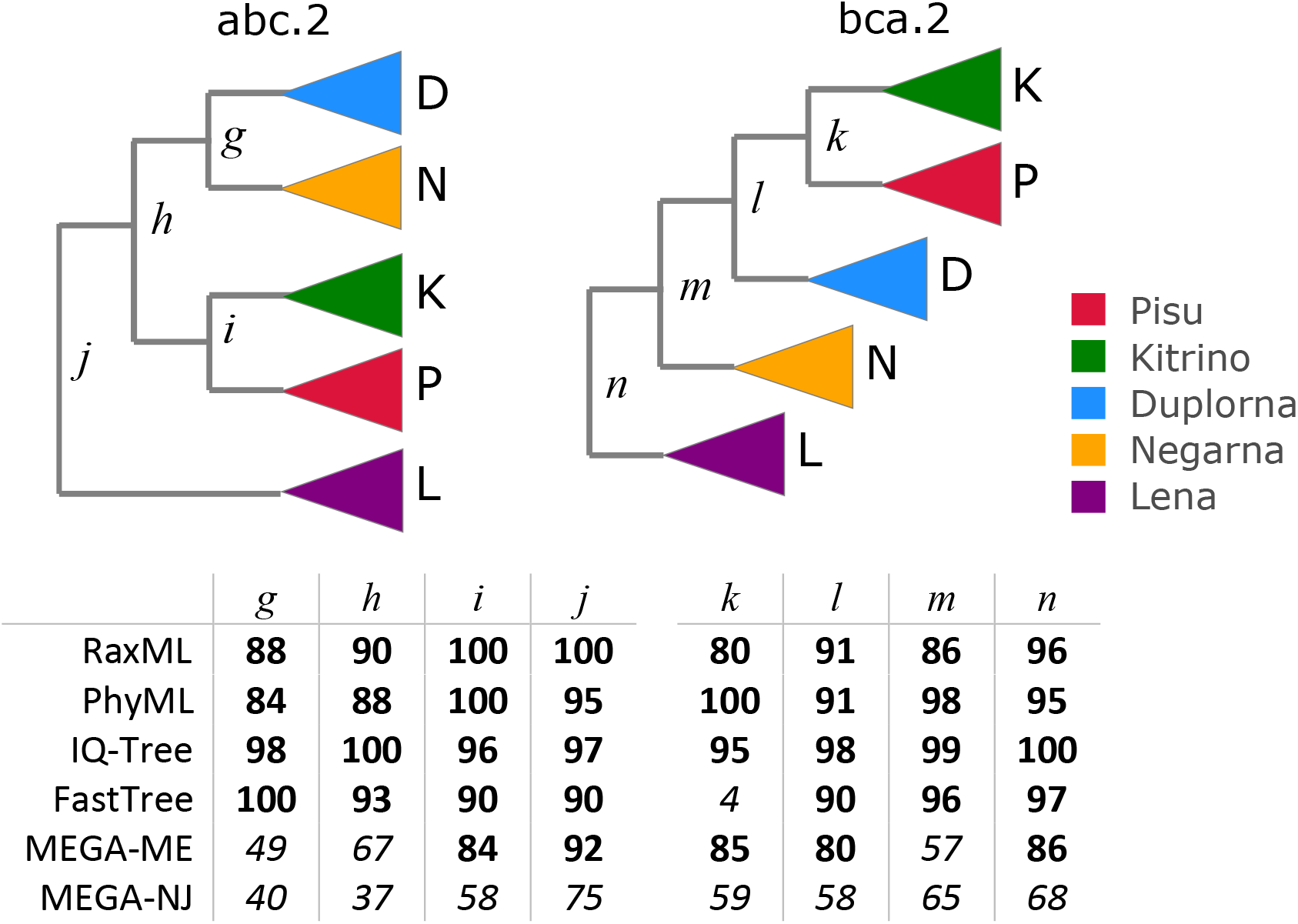
Phylum topologies estimated from replicates abc.2 and bca.2. HMM parameters are held fixed for making the MSAs (both have random seed 2) while the guide tree topology is permuted. All six tree estimation methods agree with each other on the topology on a given MSA, but the topologies are different so one or both topologies must be wrong and the reproducible wrong tree must be induced by guide tree bias as the MSA is otherwise unchanged. Most methods give high bootstraps (shown in table below the trees) for most or all of the edges.

**Figure 7:**
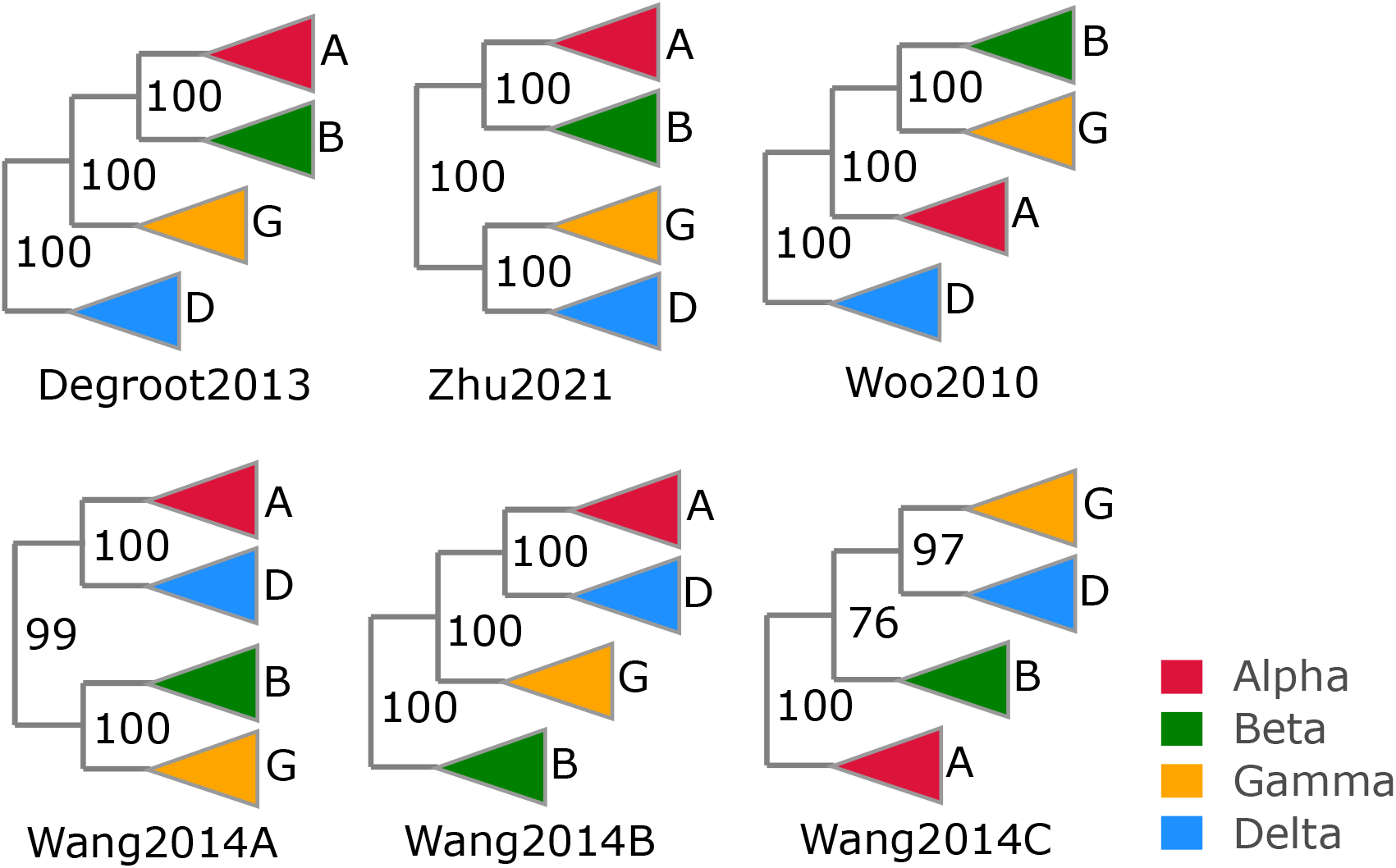
Conflicting coronavirus genus topologies in the literature. Trees from four published papers reporting high bootstraps for conflicting genus topologies: Degroot2013 [31], Wang2014 [32], Woo2010 [33] and Wang2014 [32]. Wang2014 estimated trees from three different alignments: (A) whole genomes, (B) spike protein and (C) nucleocapsid protein.

### Monophylicity of phyla in *Riboviria*

A deep RNA virus phylogeny was recently reported in [11] (hereinafter Wolf2018) and subsequently used as the basis for introducing new taxonomic ranks including phylum [17]. I measured the monophylicity of the new phyla on a diversified ensemble. For each MSA, a tree was estimated using RAxML, and the best-fit subtree identified for each phylum. To investigate whether the Wolf2018 MSA might be more accurate than Muscle5, I checked the alignment of essential catalytic residues, finding that Muscle5 is better by this measure (Fig. 9, Methods). Results are shown in Fig. 8. Panel **a** is the Wolf2018 tree topology showing the high reported bootstrap values which supported the putative monophylicity of the new phyla, panel **b** shows a typical tree from the ensemble exhibiting similarly high bootstrap values for a conflicting topology. Panel **c** shows monophyly of the phyla, where EM ranges from 41% for *Duplornaviricota* to 64% for *Negarnaviricota.* Panel **d** shows the composition of false-positive leaves under the best-fit subtree for each phylum, showing substantial mixing of all pairs of phyla (note that while *Negarnaviricota* has relatively few FPs, it has 25% FNs mixed into other subtrees). Combined, these results strongly suggest that the high bootstrap values in the Wolf2018 tree are artefacts of biases in their MSA, and the newly-adopted phyla are far from monophyletic.

**Figure 8:**
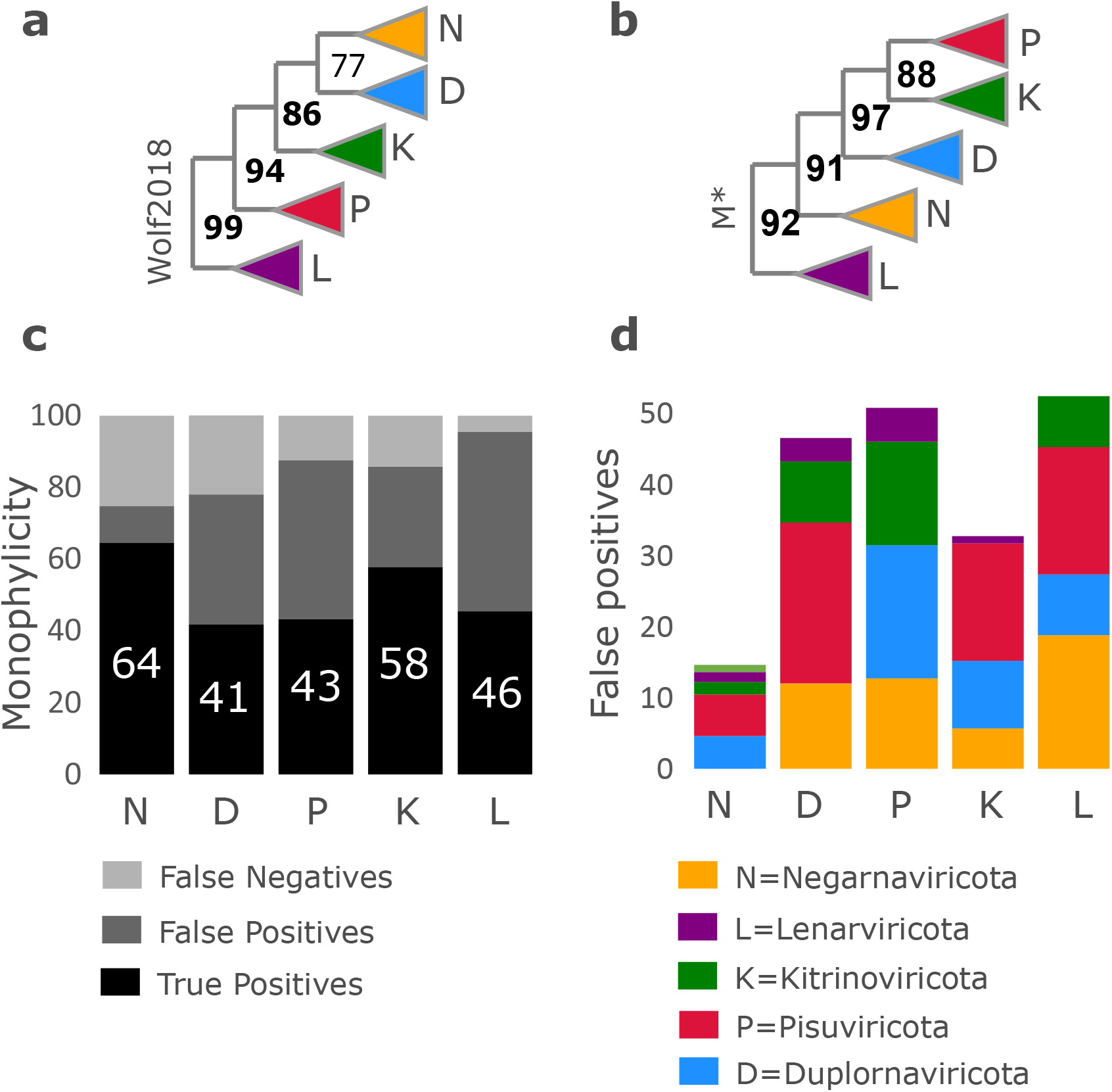
Monophylicity of RNA virus phyla. **a** Wolf2018 tree showing high bootstrap values as reported in their paper. **b** RAxML tree estimated from *M*\* (Muscle5 replicate with highest AC) showing high bootstrap values for a conflicting topology. **c** Mean false positive frequencies as a percentage of the best-fit subtree, averaged over a diverse ensemble. **d** Monophyly of the tree in panel of the best-fit subtree as percentage TP, FP and FN respectively, averaged over a diverse ensemble. Note that TP is low, ranging from 43% for *Pisuviricota* to 65% for *Negarnaviricota.* Compare with Fig. 3.**b** showing high monophyly of coronavirus genera.

**Figure 9:**
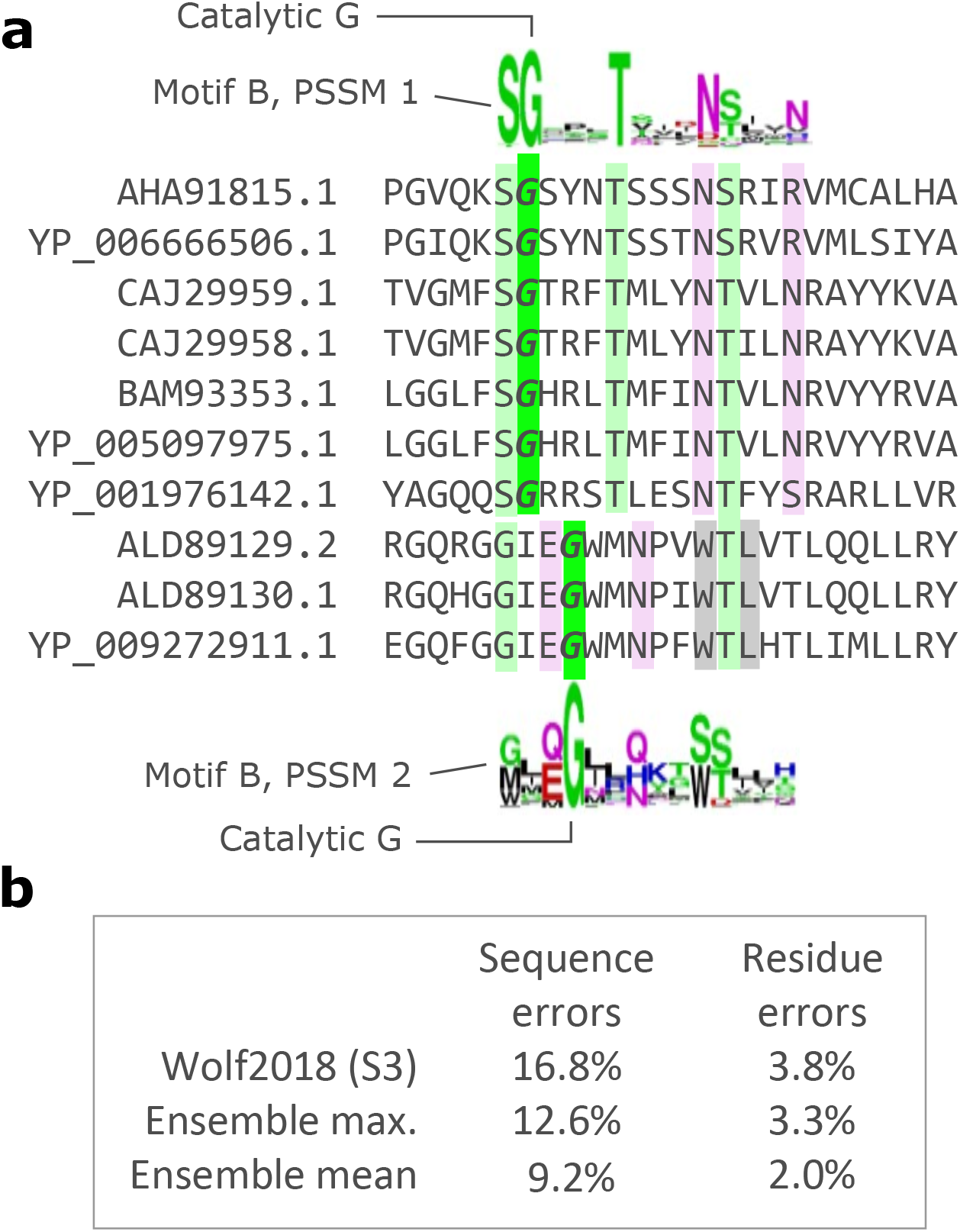
Misaligned catalytic residues in RdRp MSAs. Misalignments of essential catalytic residues were identified using `Palmscan`. **a** Ten representative sequences from the Wolf2018 alignment are shown. The top seven sequences place the catalytic glycine (G) in motif B in a different column than the bottom three. Sequence logos for the relevant `Palmscan` PSSMs are shown above and below the alignment. **b** Percentages of sequences with at least one catalytic residue misalignment and the total number of residue misalignments for Wolf2018 alignment S3 and the maximum and mean values on the corresponding `Muscle5` ensemble. S3 is a subset alignment used by Wolf2018 to estimate the top-level (near-root) branching order of their tree, it contains 238 sequences. The equivalent `Muscle5` ensemble has 249 sequences per MSA, selecting different subsets in addition to different alignment parameters to construct replicates. Note that all `Muscle5` replicates have substantially fewer errors than S3.

## Discussion

### High accuracy ensembles enable improved confidence estimates

It is *de facto* standard practice in biological sequence analysis to make a “best effort” analysis based on one preferred alignment, proceeding on the implicit assumption that this alignment is correct, or at least good enough to make the desired downstream inferences. Sometimes, columns considered to be less reliable (e.g. “gappier”) may be discarded, but the possible impact of any remaining biases are almost universally neglected, presumably due to a lack of awareness that they may be present combined with a lack of convincing methods for identifying and mitigating such errors. The results presented here show that alignment bias can have a significant impact, and as an important special case that tree bootstrapping from a single MSA may give high confidence to incorrect edges caused by bias in the MSA. In contrast to previous ensemble methods, `Muscle5` generates alignments with greater diversity of systematic errors (if present) while maintaining state-of-the-art benchmark accuracy, thereby enabling detection and mitigation of incorrect inferences due to MSA bias.

### Ensemble analysis complements bootstrapping

The Felsenstein bootstrap relies on several assumptions [18]: the alignment is correct, sites evolve independently according to Markov models, the best tree is successfully identified for each resampled alignment, and the best tree will converge on the true tree as more columns are sampled from the underlying distribution, i.e., that the evolutionary model is a good enough approximation to identify the correct tree. All these assumptions are surely violated in practice. Alignments are often wrong. Sites are not independent, and their evolution is not strictly Markovian. The space of trees is too large to search exhaustively for more than a few leaves. The best tree may not be the true tree, which seems almost certain in non-trivial cases because evolutionary models are highly simplified. Thus, bootstrapping may be unreliable, as illustrated by examples presented in this work. Ensemble analysis assumes that alignment errors are sampled sufficiently across the ensemble to induce variations in downstream inferences comparable in size with a typical inference error. This assumption is violated if the alignment is robust against parameter perturbations but nevertheless wrong. Thus, for assessment of phylogenies, bootstrapping and ensemble analysis are complementary. If there is no variation in the ensemble, then standard bootstrapping alone is appropriate. If the ensemble is variable, ensemble confidence may be more credible than bootstrapping, as shown by the examples of coronavirus genera and RNA virus phyla where bootstrapping and ensemble confidence imply different conclusions but the ensemble is more credible.

### Extending the ensemble approach

Ensemble confidence can straightforwardly be applied to other inferences from an alignment such as predicted secondary structure, paralog/ortholog discrimination and so on. The topology of putative clades can be assessed by comparing results when different subsets of sequences are selected; in particular resampling of sequences with replacement can be considered bootstrapping of rows rather than columns, noting that the rows should be realigned after resampling. If phylogeny is estimated from multiple genes, results can be compared on different subsets of genes. The Felsenstein bootstrap can be improved by sampling columns from an MSA ensemble rather than a single MSA (*ensemble bootstrap*), thereby accounting for alignment uncertainty in addition to under-sampling of columns [19]. However, I would not advocate relying on the ensemble bootstrap as a complete solution to accounting for MSA bias in phylogenetic tree estimation. While ensemble bootstrap values from a `Muscle5` ensemble will surely be more reliable than conventional bootstrapping from a single MSA, there is no substitute for intelligently assessing robustness by replicating an analysis in distinctly different ways, paying particular attention to likely sources of systematic bias.

### Reported RdRp phylogeny is not replicated

The results reported here show definitively that high bootstrap values for the RdRp-based phylogeny reported in Wolf2018 are artefacts of MSA bias. Ancient groups and their branching order are not reproduced in an ensemble analysis based on MSAs that are more accurate than the Wolf2018 alignment. This, it appears that new taxa based on these groups were introduced prematurely as they are probably far from monophyletic.

### Conclusions

Science is suffering from a replication crisis driven by practices such as p-value hacking, harking (hypothesis after result is known) and cherry-picking [20]. Sequence analysis software offers the biologist a bewildering array of alternatives for ubiquitous routine tasks such as alignment and tree-building. MUSCLE or MAFFT? Maximum likelihood or minimum evolution? RAxML or PhyML? How many discrete gamma categories should you have in your model? Common practice is to choose one protocol for a mix of stated and unstated reasons which may be more or less defensible: my colleague does it this way, it got a good benchmark score, or it gave a result I like better on my own data. Picking a single best protocol disregards the possibility that best may not be good enough. You can guard against this pitfall by performing your own replication study. This may be as simple as trying different software packages with a few different options. Even if alternative protocols are believed to be less accurate, a thoughtful comparison of the results provides a useful indication of whether the preferred protocol can be trusted. While automated sequence analysis methods should never be entirely trusted, including methods for constructing replicates, high-accuracy alignment ensembles enable a substantial improvement in assessment of inferences in many areas of molecular biology.

## Methods

### Related work

Several methods for generating collections of alternative sequence alignments have previously been described in the literature. The earliest I am aware of date from 1995 [21, 22]. In [21] the author assessed the robustness of arthropod phylogenies under variation in transversion-transition probabilities and gap penalties, noting that “[the] disturbing circularity of the interaction between the specification of [insertion-deletion and substitution probabilities] *a priori* and their inference *a posteriori* is a general and central problem in molecular phylogenetics”. The Elision method [22] concatenates variant MSAs before estimating a phylogenetic tree. A 1997 study [23] of 18S ribosomal RNA in *Apicomplexa* assessed robustness of phylogenetic inferences using multiple alignments from different software packages, finding that “different alignments produced trees that were on average more dissimilar from each other than did the different tree-building methods used”. More recent proposals along similar lines include [19, 24, 25]. `Muscle5` improves on previous ensemble methods in two crucial respects. First, all `Muscle5` replicates have high accuracy such that no other MSA is preferred *a priori*, while in previous methods there is a clearly preferred MSA, i.e. the alignment generated by the algorithm and parameter combination giving the best benchmark score. In my terminology, previous methods generated S-ensembles while `Muscle5` generates H-ensembles (Table 1). Second, `Muscle5` replicates explore a substantially greater range of possible biases by introducing more consequential variations into substitution scores and guide trees. Scaling to large datasets by a divide-and-conquer strategies was implemented in an update to `MAFFT` [26], followed by `Clustal-Omega` [27] and others. Below, the `Muscle5` algorithm is briefly described; further details of the algorithm and its improvements over previous work are provided in Supplementary Material.

### Hidden Markov model

Posterior probabilities for aligning every pair of letters in the input sequences are calculated using a hidden Markov model (HMM) with topology shown in Fig. 10. This is a coupled Markov model [10], extended to double-affine gaps as found in Probcons source code version 1.12, which differs from the HMM described in the paper [8]. For a pair of sequences *x,y,* alignment columns are emitted by the match state *M* and by insert states *I_x_, I_y_*, *J_x_* and *J_y_*. *M* emits a column containing an aligned pair of letters. *I_x_* and *J_x_* emit one letter from *x*, similarly *I_x_* and *J_x_* emit one letter from *y*. The I states induce short gaps while the *J* states induce longer gaps. Alignments begin in the start state *S*. Match state emission probabilities are obtained from the joint-probability form of the BLOSUM62 matrix [28]; insert states emit letters according to the marginal probabilities of BLOSUM62. In `Probcons`, transition probabilities were trained by expectation-maximisation on version 2 of the `Balibase` benchmark. For `Muscle5`, I chose somewhat arbitrary round numbers (Fig. 10), guided by the defaults in Probcons and the premise that small differences should be immaterial to alignment quality. Symmetries are enforced between *x* and *y* and between sequences and reversed sequences, giving five independent sets of probabilities for mutually exclusive alternative events: (1) transitions from the start state *S* (symmetric with transitions into the end state *E*), (2) transitions out of *M*, (3) transitions out of *I*, (4) transitions out of *J*, and (5) letter-pair emissions.

**Figure 10:**
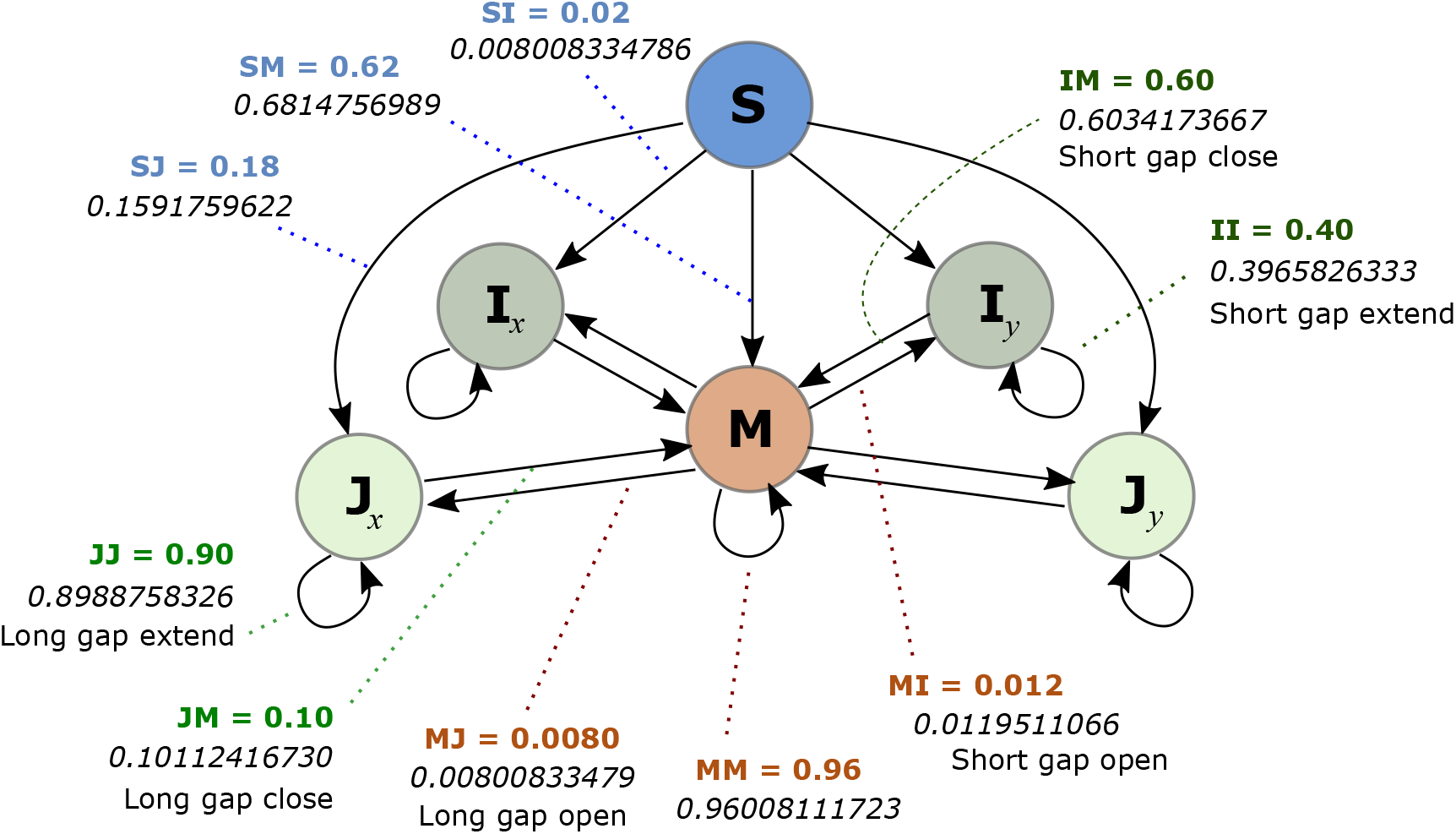
Hidden Markov model. HMM used by the `Muscle5` algorithm, showing transition probabilities, which are equivalent to gap penalties. There are two insert states (I and ***J***) for each sequence. Default probabilities for `Muscle5` are shown in bold, coloured text; defaults for `Probcons` are italic. Probcons values are specified to 10 significant figures per its source code, while `Muscle5` values are rounded to two significant figures on the premise that small differences should be immaterial to alignment quality. Emission probabilities are taken from the joint-probability form of the BLOSUM62 matrix (not shown).

### Muscle5 algorithm

The core component of `Muscle5` is a parallelised re-implementation of Probcons. Iterative consistency transformations are applied to the posterior probabilities (two rounds by default); a greedy maximum expected accuracy guide tree is constructed; and refinement by randomised partitioning is applied (100 rounds by default). Scaling to large datasets is achieved by a divide-and-conquer strategy (the `Super5` algorithm); details in Supplementary Material.

### HMM perturbations

Perturbations are introduced by adjusting all probabilities according to the rule *P* → (1 + *αδ*)*P*, where *α* (amplitude) is a constant and *δ* is a random number uniformly distributed between –1 and +1. By default, *α* = 0.25, causing perturbations up to ±25%. This value was chosen by trial and error on a subset of Balibase. To maintain normalisation (i.e., ensure that 0 < *P* < 1 for all *P* and the sum over mutually exclusive events is 1), a second adjustment *P_k_* → *P_k_*/∑*_i_P_i_* is applied after all probabilities have been perturbed by the first rule. The sum in the denominator is over probabilities in the same set of mutually exclusive events as *k*, e.g. transitions from M. A single parameter (*α*) sets a scale for all perturbations, reducing possible concerns about over-fitting (too many parameters) and over-tuning (choosing a value that performs well on the training set but poorly on new data) to a minimum.

### Guide tree permutations

The goal of permuting the guide tree is to induce substantive variation into any systematic errors due to progressive alignment, without compromising accuracy. Optimising accuracy requires that closely-related sequences are aligned before more diverged sequences are added [3, 6], which in turn requires that the guide tree joining order should be preserved close to its leaves. Substantive variations require that larger groups are joined in different orders. These constraints imply that the tree should be mostly unchanged close to the leaves and large changes should be made to the joining order of larger groups close to the root, but these goals can be conflicting in practice as guide trees are often highly unbalanced, i.e. many nodes join small groups to large groups, in which case naive re-arrangements of the tree may fail to induce substantive variations. With these considerations in mind, `Muscle5` manipulates the guide tree *T* as follows. An edge is identified which divides the leaves of T into subsets *a* and *bc* such that the ratio |*a*|/|*bc*|≈ 1/2, i.e. *a* has approximately one third of the leaves in *T*. The tree *bc* is then divided into subsets *b* and *c* of equal size so that |*b*|/|*c*| ≈ 1. Regardless of the original guide tree topology, when there are many leaves this procedure successfully divides *T* into three subtrees *a, b* and *c* of approximately equal size where the joining order close to the leaves is mostly preserved. Progressive alignment is performed using the original guide tree and permutations ((*a, b*), *c*), ((*a, c*), *b*) and ((*b,c*),*a*), abbreviated to *none, abc, acb* and *bca* respectively. A replicate is identified as *perm.s,* e.g. abc.3, where perm is the guide tree permutation and *s* is the random number seed, where the special case *s* = 0 indicates that no perturbations are applied, i.e. default HMM parameters are used. See Supplementary Materials for further details and discussion.

### Diversified ensemble

A diversified ensemble is designed to maximise variation among replicates by setting the random number seed *s* = 0,1…*N* where *N* is the desired number of replicates, while guide tree permutations cycle through the four variants *none, abc, acb* and *bca*. For the results reported in this work, *N* = 100 was used. Alternatively, convergence criteria could be set which terminate generation of further replicates when sufficient diversity has been sampled, though this feature was not implemented in the code described here. For convergence, I suggest an upper limit of *m* replicates where *m* is the median number of columns in the ensemble so far, and also testing the number of singleton distinct columns (*n*_1_, found in exactly one replicate) compared to the number of reproduced distinct columns (*n*_2_, found in two replicates). If *n*_2_ > *n*_1_, most of the potential diversity in the ensemble has been sampled. In practice, when sequences are closely related most of all replicates may be identical; in such cases setting convergence criteria would save substantial computing resources in high-throughput applications.

### Best-fit subtree

Given a tree and categories assigned to a subset of its leaves (e.g., phylum names), the best fit for category *C* is identified as the node *t* which minimises the number of errors in its subtree. The number of true positives (TP) is the number of leaves under *t* which belong to *C*. Errors include false positives (FP, i.e. leaves under *t* which are not in *C*), and false negatives (FN, i.e. leaves in *C* which are not under *t*).

### Ensemble Monophyly

Given a tree and a category *C* (e.g. phylum name), the best-fit subtree *t* is identified. Monophyly of *C* is then *m* = *TP*/(*TP* + *FP* + *FN*). If *C* is monophyletic, *m* =1, otherwise *m* < 1, and m is smaller with increasing errors. Ensemble Monophyly is mean *m* over an ensemble of trees.

### Root identification by an out-group

Given a tree and an out-group category *C*, the best-fit subtree is identified after assigning all leaves not belonging to *C* to a second category. The root is then placed in the edge joining the best-fit subtree of *C* to the rest of the tree. This procedure accommodates the trivial case where *C* is monophyletic in the estimated tree, and also more difficult cases where *C* is polyphyletic.

### Condensed tree

Given a rooted tree *T* and category labels on a subset of its leaves (e.g., phylum names), the best-fit node is identified for each category. Edges in the path from each best-fit node to the root are preserved, all other edges are deleted. This produces a tree *T_b_* in which all leaves are best-fit nodes. Each unary node *u* in *T_b_* is collapsed by replacing *u* and its incoming and outgoing edges *u_i_* and *u_o_* by a single edge *e_u_*; this is repeated until all nodes have degree > 1. The length of *e_u_* is the sum of the lengths of *u_i_* and *u_o_*; the bootstrap value assigned to *e_u_* is the larger of the bootstraps for *u_i_* and *u_o_*. The resulting tree is the condensed tree of *T* according to the category labels. A condensed tree summarises the branching order of its categories, assuming they are monophyletic or approximately monophyletic so that best-fit nodes are estimates of most recent common ancestors.

### RdRp alignment quality

The `Palmscan` algorithm [29] uses position-specific scoring matrices (PSSMs) to identify three well-conserved motifs in the RdRp palm domain which are conventionally designated *A, B* and *C* respectively. These motifs include six essential catalytic residues: two aspartic acids (`D`) in motif *A*, a glycine (`G`) in motif *B*, and `GDD` in motif *C* [30]. The quality of an RdRp alignment was assessed by using `Palmscan` to identify the position of each of these catalytic residues in all sequences which successfully matched the PSSMs. If a catalytic residue appears in a different column from the majority of other sequences, it is considered to be misaligned (Fig. 9). The total number of misaligned catalytic residues was used as a quality metric.

### Validation on simulated data

Alignment and tree inference methods are often validated by simulating sequence evolution *in silico*. I chose not to do so in this work. Simulations employ drastically simplified models of evolution which are similar to the drastically simplified models used by multiple alignment and tree inference algorithms. In reality, sequence evolution is an enormously complex process disrupted by historical contingencies ranging from fortuitous outcomes of DNA repair machinery failures and narrowly-won host-pathogen arms races to asteroid impacts. Therefore, simulations of deep evolutionary history are at best suggestive and at worst entirely uninformative if one is interested in real biology. Simulations also exacerbate a common sociological problem in computational biology, namely that the developers of a new method have an opportunity to fish for significance before publication. Confronted with disappointing results, authors may rationalise tweaking a simulation until improved (simulated) performance is obtained for their method. These considerations beg the question of whether simulations could convincingly support the main claims of this paper, which are (1) `Muscle5` MSA replicates have high and practically indistinguishable accuracy, and (2) the effects of alignment errors can be assessed by comparing inferences from different replicates. Claim (1) is supported by results on structure-based benchmarks. While structural similarity does not necessarily imply sequence homology, structural alignments are largely independent of sequence, and greater agreement with structural alignments therefore surely correlates strongly with more accurate alignment of homologous residues. If a simulation fails to recapitulate relative algorithm performance according to structure, the failure is more plausibly explained by a defect in the simulation than a defect in the structural benchmark. Claim (2) is self-evidently true because two different alignments of the same sequences cannot both be correct.

## Software availability

Binaries and source code are available under the GPL v3 license at: https://github.com/rcedgar/muscle

https://github.com/rcedgar/newick

Code and data required to reproduce experimental results are available under the GPL v3 license at:

https://github.com/rcedgar/rdrp_tree_experiments

## Software commands

Software versions and command lines are provided in Supplementary Material.

## Acknowledgements

I am grateful to Alexandros Stamatakis and Artem Babaian for helpful discussions. Some of the computing resources used in this work were kindly provided by the University of British Columbia Community Health and Wellbeing Cloud Innovation Centre, powered by AWS.

